# Optogenetic olfactory behavior depends on illumination characteristics

**DOI:** 10.1101/559674

**Authors:** Tayfun Tumkaya, James Stewart, Safwan B. Burhanudin, Adam Claridge-Chang

**Affiliations:** Institute for Molecular and Cell Biology, A*STAR, Singapore 138673; Department of Physiology, National University of Singapore, Singapore; Program in Neuroscience and Behavioral Disorders, Duke-NUS Medical School, Singapore

## Abstract

Optogenetics has become an important tool for the study of behavior, enabling neuroscientists to infer causations by examining behavior after activating genetically circumscribed neurons with light. Light-induced neural activity is affected by illumination parameters used in experiments, such as intensity, duration, and frequency. Here, we hypothesized that the intensity of light and the presence of oscillations in illumination would alter optogenetically induced olfactory behaviours. To test this, we activated olfactory receptor neurons (ORNs) in *Drosophila* by using either static or pulsed light stimuli across a range of light intensities. The various regimes elicited distinct behavioral valence responses (attraction, aversion, indifference) from several ORN types. Our results demonstrate the importance of both frequency and intensity for interpreting optogenetic behavioral experiments accurately; successfully generalizing optogenetic results requires the use of more than a single illumination regime.

## Introduction

For scientific research, interventional methods have been central to establishing causal relationships (Lewin, 1947; Midgley, 2003; Miesenböck & Kevrekidis, 2005; Thiese, 2014). Genetic lesion studies, for example, are critical to determine gene function (Baratz et al., 2010; Chong et al., 2015; Katsanis, 2016; Muenke et al., 1994). Similarly, neuronal activation and inhibition experiments identify links between behaviours and neurons (Bernstein & Boyden, 2011; Deisseroth, 2015; Fenno, Yizhar, & Deisseroth, 2011; Packer, Roska, & Häusser, 2013). For analyzing the function of neural circuits, optogenetic activation has become one of the most-used techniques.

Using light to actuate photosensitive proteins that are encoded by DNA enables neuroscientists to rapidly manipulate genetically circumscribed neurons (Miesenböck, 2009). Following pioneering methods used to control neuronal function with light (Banghart, Borges, Isacoff, Trauner, & Kramer, 2004; Lima & Miesenböck, 2005; Szobota et al., 2007; Volgraf et al., 2006; Zemelman, Lee, Ng, & Miesenböck, 2002; Zemelman, Nesnas, Lee, & Miesenbock, 2003), the microbial opsins have found the widest application in optogenetics (Deisseroth, 2015; Fenno et al., 2011). Microbial opsins are photosensitive seven-transmembrane proteins, typically ion pumps, channel proteins, or regulators of ion channels. Upon light activation, they modulate transmembrane voltage (Fenno et al., 2011; Miesenböck, 2009). The blue-light sensitive cation channel Channelrhodopsin-2 (Chr2) was the first microbial opsin used to control neuronal activity (Boyden, Zhang, Bamberg, Nagel, & Deisseroth, 2005). Chr2 has been iteratively engineered for improved photoconductivity, as well as for diverse wavelength sensitivities (Deisseroth, 2010; Guru, Post, Ho, & Warden, 2015; Lin, 2011). A multitude of the Chr2 variants have been used *in vivo* in animal models—such as *Drosophila*, zebrafish, *C. elegans*, and mice—to activate neurons and study behavior (Fenno et al., 2011).

The action potential of an optogenetically activated neuron is mainly determined by biophysical properties of the opsin, as well as characteristics of the light stimulus: pulse intensity, width, duration, and frequency (Boyden et al., 2005; Miesenböck, 2011; Mohanty & Lakshminarayananan, 2015). Ideally, illumination protocols are optimized to make artificial action potentials similar to the natural activity (Allen, Singer, & Boyden, 2015; Packer et al., 2013). However, there are no overall guidelines for picking light parameters for optogenetics experiments; as a consequence, illumination protocols vary greatly across studies (Herman, Huang, Murphey, Garcia, & Arenkiel, 2014; Mattis et al., 2011). How does the light-stimulus variability affect optogenetically induced behaviours?

Here, we addressed this question with the optogenetic actuation of *Drosophila* olfactory behaviour. Flies may produce directed locomotion responses (attraction or aversion) to optogenetic activation of single classes of olfactory receptor neurons (ORNs) (Bell & Wilson, 2016). In a custom-built behavioral assay, we tested the attraction or aversion valence to optogenetic activation of four classes of olfactory neurons (the ORNs expressing Or59c, Or85c, Or92a and the CO_2_-sensitive gustatory receptor neuron expressing Gr21a) by using either static or pulsed light stimulus at three different intensities. In all four cases, the valence responses to pulsed light were dramatically different to that of static light. Moreover, olfactory responses considerably varied across the different light intensities. Despite using the same opsin and behavioral assay to activate the exact same set of neurons, different light intensities and frequencies gave rise to distinct conclusions. These results reveal that the temporal structure of optogenetic light has a large impact of behavioral outcomes, and suggests that experiments using a single parameter may not be generalizable.

## Material and methods

### Fly care and strains

Flies were prepared as previously described (Mohammad et al., 2017). Briefly, flies were kept in the dark on a regular fly medium (Temasek Life Sciences Laboratories, 2018), and transferred on 0.5 mM all-trans-retinal (Sigma-Aldrich, USA) supplemented food two days prior to optogenetic experiments. Cantonized *w*^*1118*^ flies were used as wild type. The *ORN-Gal4 (Couto, Alenius, & Dickson, 2005; Fishilevich & Vosshall, 2005)* and *UAS-CsChrimson (Klapoetke et al., 2014)* stocks were obtained from the Bloomington Drosophila Stock Center. All flies used in the study were starved on 2% agarose for 12–18 hours prior to the experiments.

### Optogenetic behavior apparatus

An optogenetic apparatus was designed somewhat similar to one previously described (Mohammad et al., 2017). A rectangular arena (11.5 × 14.5 × 0.3 cm) was cut out of an acrylic board with 16 tracks (50 × 4 mm) carved on it to place flies. The arena was kept in an incubator to maintain a dark and 25°C degrees environment during experiments. Six red LEDs (617 nm; Luxeon Rebel LEDs on a SinkPAD-II 10mm Square Base) glued on heat sinks were placed above the arena at a ∼45 angle on both sides. The LEDs were equipped with optic lenses (17.7° 10mm Circular Beam Optic) and powered with 700-mA BuckPuck drivers. With the aid of masks cut from black acrylic, separate control of the LEDs on each side allowed to illuminate red light on either half of the arena. The red LEDs were controlled by custom software, CRITTA (Mohammad et al., 2016). The light intensity was measured by using a thermal power sensor (Thorlabs S310C) connected to a power and energy-meter console (Thorlabs PM100D). Two grids of infrared LEDs illuminated the platform for an AVT Guppy PRO F046B camera, which was equipped with an IR bandpass filter (Edmund Optics, Singapore), to monitor flies. The camera streamed video to the CRITTA software to detect fly positions in real-time. The fly-position data were logged in text files for offline analysis.

### Optogenetic Self-Administration Response (OSAR) assays

Flies were collected 2-3 days prior to the experiments. They were loaded into the chambers after being anesthetized by a 30-sec ice treatment. The experimental procedure consisted of acclimatization of flies for 30 s, illumination of left half of the arena for 45 s, a recess of 30 s, illumination of right half of the arena for 45 s, and a final interval of 30 s to ensure a consistent breaks between the experiments. Each group of flies was tested at three intensities—4.65, 9.3, 14 µW/mm^2^–either in static or pulsed (20 Hz, 50% duty cycle) light. The frequency for pulsed light was chosen following a previously published method (Bell & Wilson, 2016), and the pulsed light was normalized to have the same intensity as the static light.

### OSAR data analysis: wTSALE metric

Fly position data recorded by the CRITTA software were analyzed by custom-built Python scripts. The light preference of flies is determined by analyzing the proportion of time they spend in the illuminated zone once they discovered it. We termed this metric Time Spent After Light Encounter (TSALE). Given that some flies discover the light from the start, and some never do, we weighted TSALE linearly with a number ranging from 0 to 1: flies that never encountered the light could not inform us about the light preference and had a weight of 0; flies who discovered the light from the start provided the most insight and weighted by 1 (weighted-TSALE; wTSALE). The wTSALE score was calculated for each fly, and then averaged for control and test genotypes. The mean difference between the control and test groups was taken as the effect size (ΔwTSALE); mean difference distributions and 95% bootstrapped confidence intervals (CIs) were calculated by using the DABEST Python package (Ho, Tumkaya, Aryal, Choi, & Claridge-Chang, 2018), and presented in the following format: ‘*mean* [95CI *lower bound, upper bound*].’

### Data and code availability

The Python scripts used to analyze fly tracking data are available on GitHub at *‘https://github.com/ttumkaya/WALiSuite_V2.0’*, as well as on the Zenodo server at *‘zenodo.2540290’*. All the tracking data can be found on the Zenodo server at *‘zenodo.2541462’*.

## Results

### Triggering valence with optogenetic activation

Artificial activation of single classes of *Drosophila* sensory neurons has been shown to be able to drive valent behavior: attraction or aversion, depending on the neuronal type (Bell & Wilson, 2016; Hernandez-Nunez et al., 2015; Suh et al., 2007). To examine the valent behavior elicited by olfactory-neuron activation, we used a behavioral assay in which flies can be given a choice between red light or no light; we refer to it as the Optogenetic Self-Administration Response (OSAR) assay (Fig 1A). Transgenic flies that express a red-light-sensitive cation channel, CsChrimson (Chr), in neurons of interest are loaded in the OSAR arena. Flies can either spend increased time in the red light, avoid it, or display indifferent behavior. We calculated the relative valence of activation by measuring how much time flies spent in red light after they first encountered it (Fig 1B). To test the hypothesis that the characteristics of optogenetic light stimulus influence the behaviour, we used either pulsed light at 20 Hz (Bell & Wilson, 2016) or static illumination at three light intensities.

**Figure 1.**
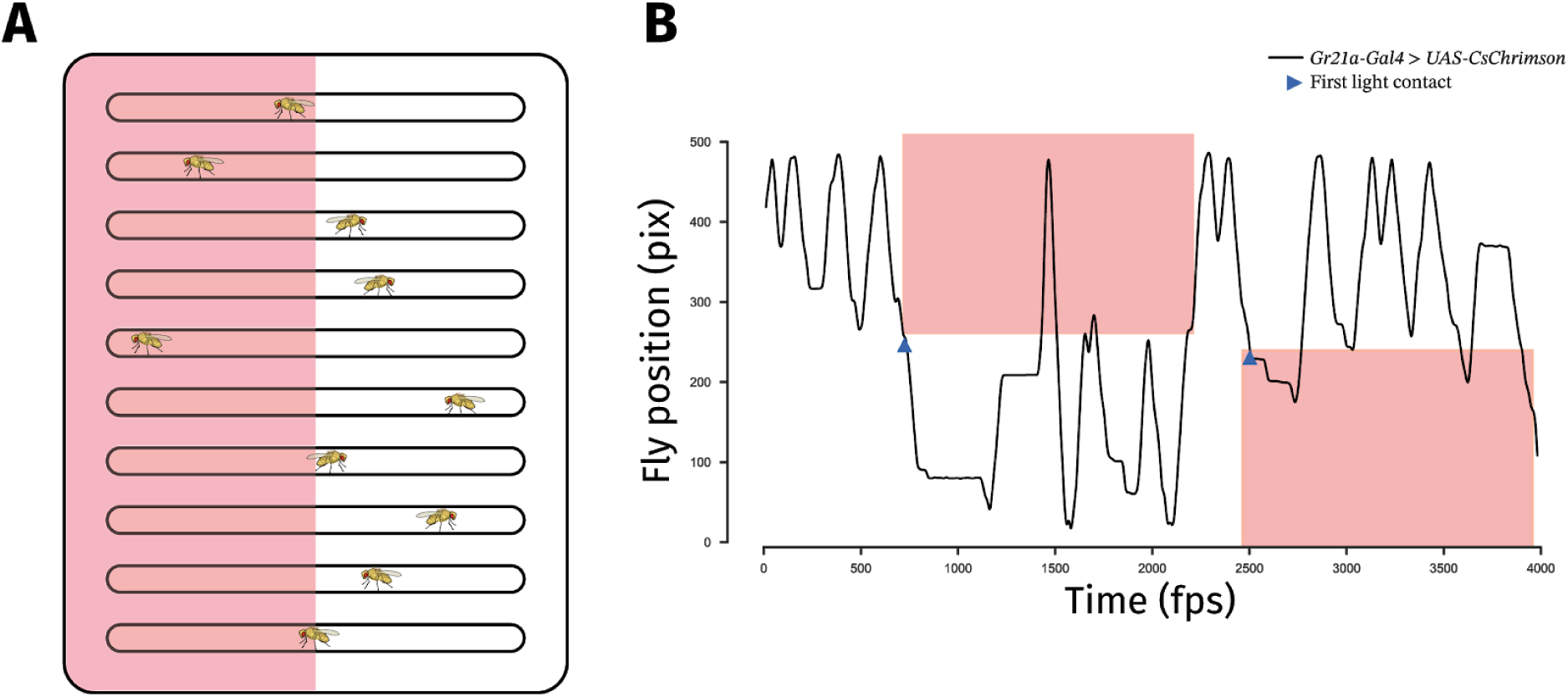
The Optogenetic Self-Administration Response (OSAR) assay. **A.** Flies are loaded in a darkened arena and given a choice between red light and dark. **B.** An example trace of a fly that spends most of its time in the dark zones after encountering red light (blue arrows). Red boxes indicate when the fly is walking through red illumination.

### Flies avoid pulsed activation of Gr21a neurons

For *Drosophila*, CO_2_ acts a stress odor and triggers an avoidance response (Suh et al., 2004). The gustatory receptor Gr21a is one of the two receptors that mediate CO_2_ aversion (Jones, Cayirlioglu, Kadow, & Vosshall, 2007; Kwon, Dahanukar, Weiss, & Carlson, 2007; Suh et al., 2004). Artificial activation of Gr21a neurons is sufficient to elicit avoidance (Suh et al., 2007). Here, we tested Gr21a avoidance by optogenetically activating these neurons with either static or pulsed light. While continuous photoactivation was aversive only at the lowest light intensity (4.65 μW/mm^2^, Fig 2A), pulsed light triggered aversion at all three intensities (Fig 2B). This dramatic behavioral difference between static- and pulsed-light activations indicates that the presence or absence of oscillations can alter the behavioral effect of optogenetic actuation (Fig 2C).

**Figure 2.**
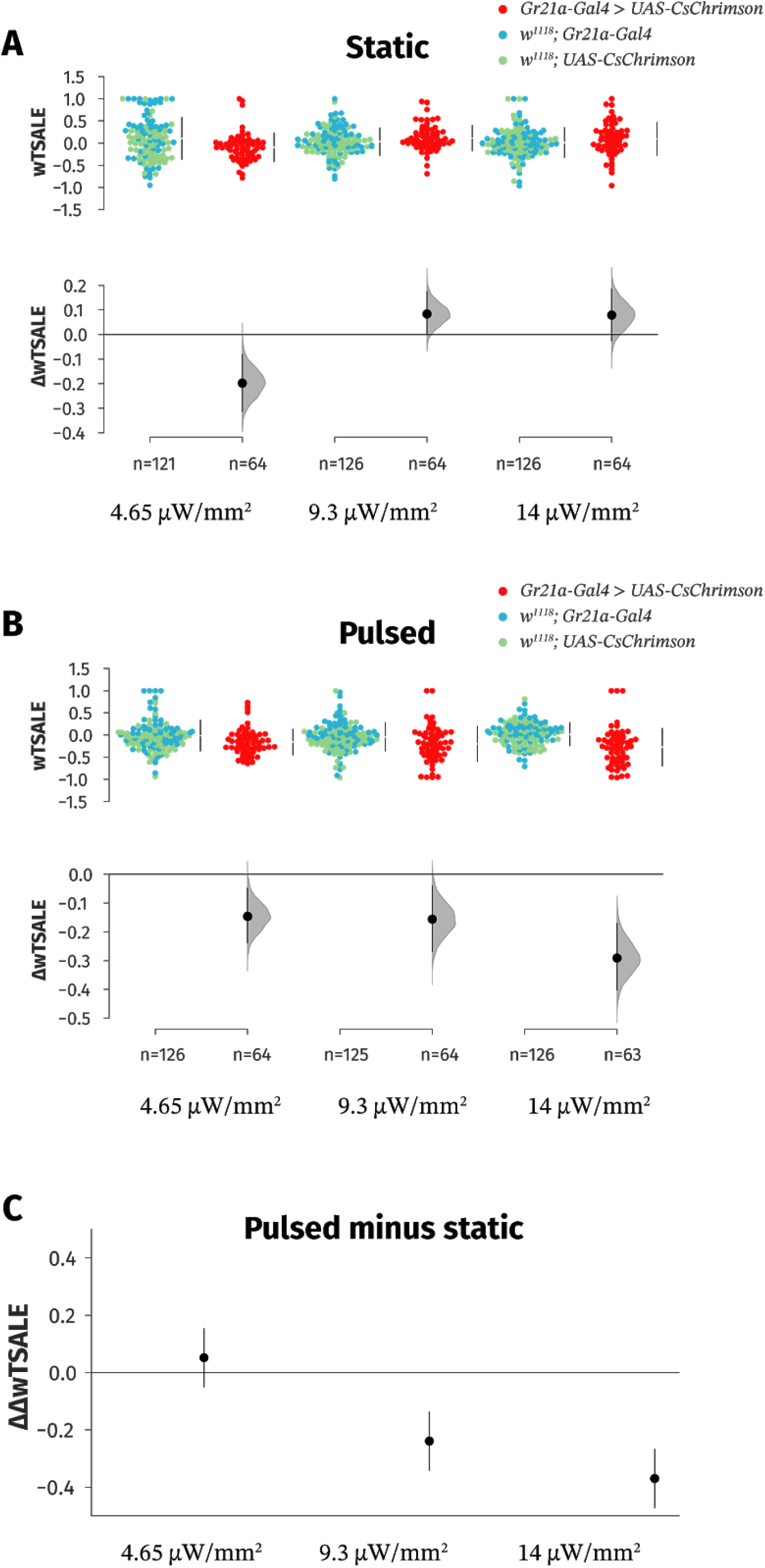
Pulsed-light and static-light stimulation of the Gr21a neurons triggers distinct responses. **A.** In static light, flies bearing the red-light-shifted opsin Chr in the CO2-sensitive Gr21a neurons spent less time in red light only at the lowest light intensity (4.65 μW/mm^2^) relative to the parental controls. Upper plot illustrates the amount of time flies spent in red light, where the dots represent single flies. Test flies are shown in red (*Gr21a-Gal4 > UAS-CsChrimson*) and controls shown in blue and green, respectively (*w*^*1118*^; *Gr21a-Gal4* and *w*^*1118*^;*UAS-CsChrimson*). Lower plot indicates the mean wTSALE difference. All error bars are 95% CIs. **B.** When pulsed light used to stimulate the Gr21a neurons, test flies avoided red-light illuminated zones at all three light intensities (4.65, 9.3, 14 μW/mm^2^) compared to parental controls. **C.** The differences between static- and pulsed-light regimes show that pulsed stimulation of the Gr21a neurons elicits stronger aversion in the two higher light intensities (9.3, 14 μW/mm^2^): −0.24 [95CI −0.14, −0.34] and −0.37 [95CI −0.48, −0.26], respectively.

### Static-light activation of vinegar-sensing neuron evokes attraction

Unsurprisingly, vinegar flies are strongly attracted to vinegar. Vinegar has been shown to activate six glomeruli in the *Drosophila* brain, two of which are sufficient to produce the attraction behavior: Or42b and Or92a (Semmelhack & Wang, 2009). Static- and pulsed-light optogenetic activations of Or92a induced different behaviors: static-light activation became increasingly attractive in the two higher light intensities (9.3, 14 μW/mm^2^, Fig 3A), whereas pulsed light did not trigger any behavioral response (Fig 3B). The discrepancy between the two types of light stimuli is largest at the highest light intensity with a ΔwTSALE of −0.22 [95CI −0.35, −0.09] (Fig 3C).

**Figure 3.**
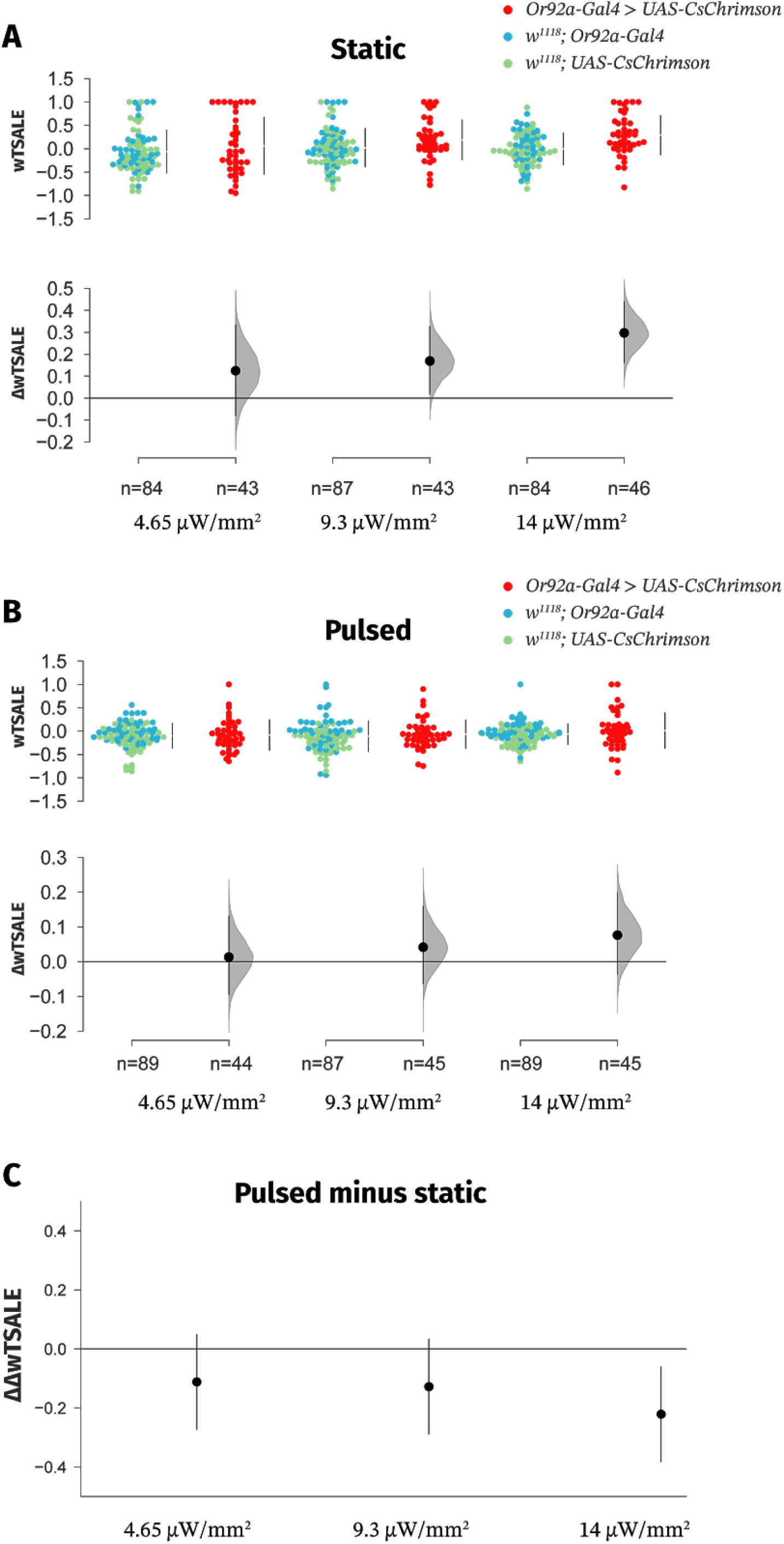
Only static illumination of the vinegar-sensitive neuron Or92a triggers attraction. **A.** Static-light stimulation of the Or92a neurons is attractive to the flies at intensities of 9.3 and 14 μW/mm^2^. **B.** Activating the Or92a neurons with pulsed light elicits only minor valence responses. **C.** The amount of time spent in red light decreases by −0.22 [95CI −0.35, −0.09] in the highest light intensity when pulsed-light is used.

### Static- and pulsed-light activations of Or59c and Or85c elicit distinct valence responses

Of the ∼60 types odor receptors expressed in the fly head, innate valence properties have been tested for a subset (Bell & Wilson, 2016). We tested two receptors (Or59c and Or85c) with unknown valence responses. Stimulation of the Or59c receptor neurons with static light produced strong aversion at the lowest light intensity (Fig 4A); the effect was markedly lower in the pulsed-light stimulation (Fig 4B-C). For the Or85c receptor neurons, while static light did not elicit any valence (Fig 4D), pulsed-light activation evoked aversion in the two lower light intensities (Fig 4E-F). These results further validated that static- and pulsed-light stimuli may elicit considerably different behavioral responses.

**Figure 4.**
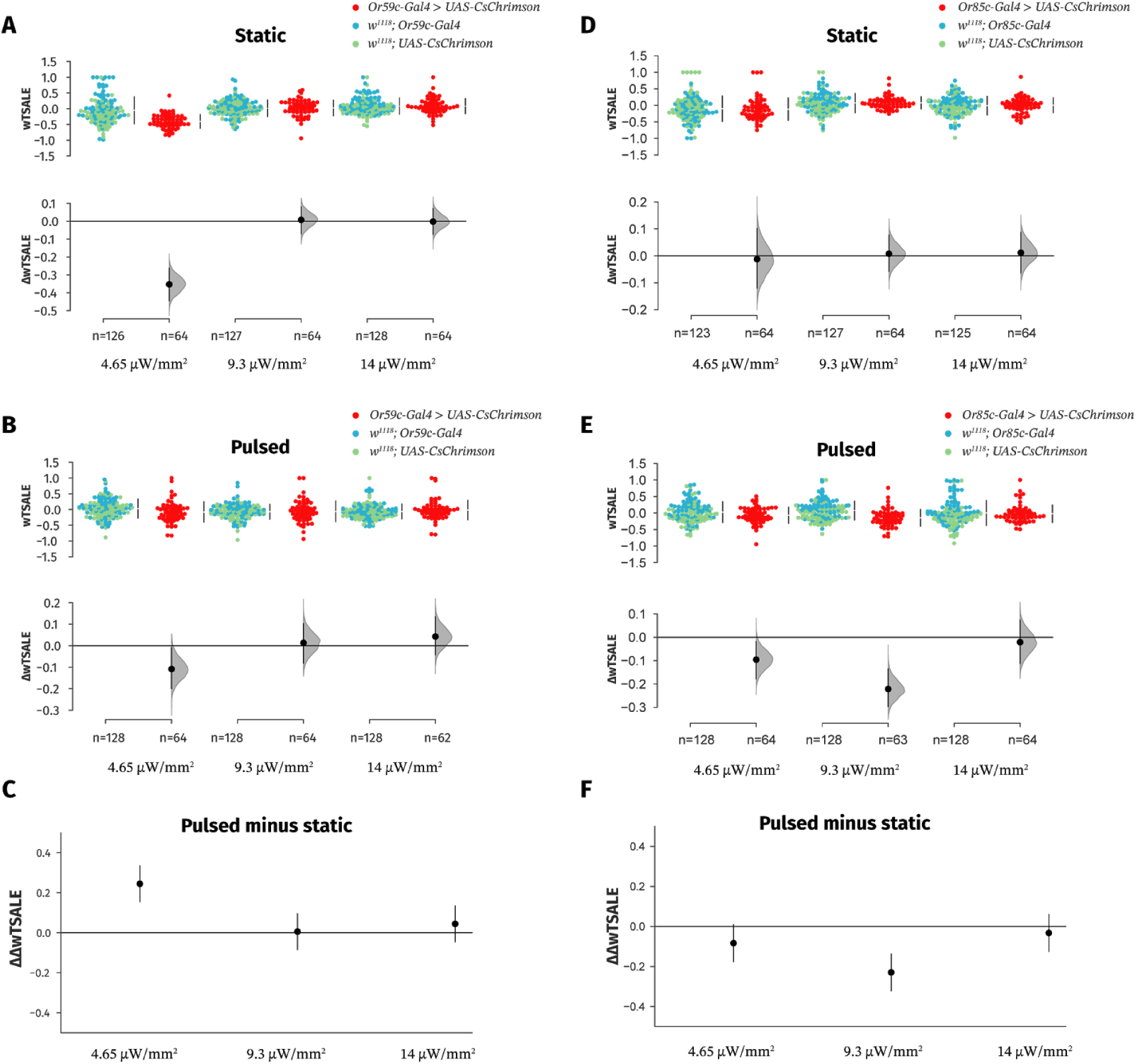
Static and pulsed light stimulations evoke different valence responses in Or59c and Or85c neurons. **A.** Static-light stimulation of the Or59c neurons is strongly aversive at the lowest intensity 4.65 μW/mm^2^. **B.** Pulsed-light stimulation of the Or59c neurons is aversive at the lowest intensity 4.65 μW/mm^2^. **C.** Pulsing the light stimulus reduced the Or59c-induced aversion by a wTSALE of 0.24 [95CI 0.15, 0.34] at 4.65 μW/mm^2^. **D.** Static-light stimulation of the Or85c neurons does not elicit any valence response. **E.** Pulsed-light stimulation of the Or85c neurons is aversive at the two lower intensities. **F.** Static and pulsed-light stimulations of the Or85c neurons elicit substantially different valences at two lower light intensities: −0.08 [95CI −0.18, 0.00] and −0.23 [95CI −0.30, −0.16], respectively.

## Discussion

### Optogenetic temporal structure influences olfactory-related behavior

Vinegar and CO_2_ are important cues for *Drosophila* to find food and sense stress (Jones et al., 2007; Semmelhack & Wang, 2009; Suh et al., 2004). Optogenetic activation of vinegar- and CO_2_-sensing cells (the Or92a and Gr21a neurons, respectively) reproduced the natural approach and avoidance responses triggered by their corresponding ligands. However, the temporal dynamics that elicited ligand-type responses were different for the two sets of neurons: Gr21a-neuron aversion occured in response to pulsed light, while vinegar-responsive Or92a neurons drove attraction in response to static illumination. A simple explanation of this disparity is that the olfactory system uses both ORN identity and rate to drive behavior (van Breugel, Huda, & Dickinson, 2018). The Gr21a neurons have been shown to respond to rapid changes in CO_2_ concentration levels, while vinegar-sensitive neurons encode intermittent signals poorly (Faucher, Hilker, & de Bruyne, 2013). Thus, even though the pulse frequency is similar to the action-potential frequencies observed during odor responses (Bell & Wilson, 2016), it seems likely that pulsed-light stimulation of the Or92a neurons generated activity patterns are distinct from vinegar exposure.

Static- and pulsed-light stimulations induced distinct behaviours through Or59c and Or85c neurons as well: static light triggered the strongest avoidance response from the Or59c neurons; conversely, only pulsed light elicited an avoidance response from the Or85c neurons, but not static light (Fig 4A-F). Interestingly, all of the olfactory responses induced by Or59c and Or85c neurons occured at the lower light intensities (Fig 4A-B-E), and did not increase as the light intensity increased.

Of the four ORNs tested in the present study, static-light stimulation performed better in eliciting olfactory responses than the pulsed light for two ORNs (Or92a, Or59c); and the opposite was true for the other two ORNs (Gr21a, Or85c). These results suggest that neither stimulation type is necessarily superior to the other: static- or pulsed-light stimulation can capture more of the native responses than the other in inducing olfactory behavior, depending on the neuronal type.

### Temporal structure is relevant to optogenetic input in a range of circuits

ORNs are not the only neurons that drive different behaviors in response to different optogenetic activation patterns (Clark et al., 2018). An examination of crawling behaviour in the *Drosophila* larvae used activation of motor neurons with either Channelrhodopsin-2 (Chr2) or a mutant variant of it, Chr2^H134R^, that has increased responsiveness to light stimulation (Nagel et al., 2005; Pulver, Pashkovski, Hornstein, Garrity, & Griffith, 2009). Static or pulsed light was used to actuate the opsins: neither affected the behaviour when the motor neurons express Chr2; as for the Chr2^H134R^ variant, only static light affected crawling behaviour, and not pulsed light. Two recent studies found discrepant effects on sleep when the galanin neurons in the mouse ventrolateral preoptic area were activated at different frequencies: activation at higher frequencies increased wakefulness (Chung et al., 2017), while lower frequencies (0.5-4 Hz) promoted sleep (Kroeger et al., 2018). Electrophysiology recordings have consistently shown that various light-stimulus parameters have various effects on optogenetically targeted neurons (Boyden et al., 2005; Klapoetke et al., 2014; Mattis et al., 2011; Pulver et al., 2009), and growing evidence now shows that light parameters also change optogenetically induced behaviours.

### Should optogenetic activation mimic natural activity?

The extent to which optogenetic activation needs to mimic natural neuronal activity to recreate natural behaviors is not clear (Malyshev, Goz, LoTurco, & Volgushev, 2015; Miesenböck, 2009). Unstructured optogenetic activation has successfully elicited complex behaviours in model animals: courtship behaviour (Clyne & Miesenböck, 2008), olfactory learning in flies (Claridge-Chang et al., 2009); swimming behaviour in zebrafish (Wyart et al., 2009); awakening (Adamantidis, Zhang, Aravanis, Deisseroth, & de Lecea, 2007) and learning in mice (Huber et al., 2008). On the other hand, some behaviours, such as odor information processing in zebrafish (Blumhagen et al., 2011) and in mice (Haddad et al., 2013; Smear, Shusterman, O’Connor, Bozza, & Rinberg, 2011), have been shown to be highly dependent on temporal structure. Mounting evidence, including the present study, indicates that different photostimulation regimes produce distinct behaviours (Chung et al., 2017; Kroeger et al., 2018; Pulver et al., 2009). Overall, our findings support the idea that, to inform generalizable conclusions about a neural system, a single optogenetic regime will often be inadequate. As there is no one-size-fits-all guide for choosing light parameters, using multiple intensities and frequencies will be prudent when characterizing systems with optogenetic activation.

## Conclusions

Optogenetics has been extensively used to study behavior; however, the extent to which illumination protocols influence experiment results has not been addressed thoroughly. In the present study, we investigated the hypothesis that different light frequencies and intensities would induce distinct olfactory behaviours. Our results support this hypothesis: same ORNs drove different valence responses to static- or pulsed-light activations; notably, none of the ORNs triggered contrasting valence responses (attraction vs. avoidance), but rather inconsistent results (valent vs. non-valent). We propose that manifold illumination regimes must be utilized to obtain generalizable results in behavioral studies.

## Author contributions

Conceptualization: TT, ACC; Methodology: TT, ACC; Experiments: TT, SBB; Software: TT (Python), JS (LabView); Data Analysis: TT; Hardware: TT, JS; Writing – Original Draft: TT; Writing – Revision: TT, ACC; Visualization: TT; Supervision: ACC; Project Administration: ACC; Funding Acquisition: ACC.

## Acknowledgements

The authors would like to thank Dr. Sangyu Xu for discussions on the manuscript.

